# Development of a robotic cluster for automated and scalable cell therapy manufacturing

**DOI:** 10.1101/2023.12.21.572854

**Authors:** Alice Melocchi, Brigitte Schmittlein, Alexis L. Jones, Yasmine Ainane, Ali Rizvi, Darius Chan, Elaine Dickey, Kelsey Pool, Kenny Harsono, Dorothy Szymkiewicz, Umberto Scarfogliero, Varun Bhatia, Amlesh Sivanantham, Nadia Kreciglowa, Allison Hunter, Miguel Gomez, Adrian Tanner, Marco Uboldi, Arpit Batish, Joanna Balcerek, Mariella Kutova-Stoilova, Sreenivasan Paruthiyil, Luis A. Acevedo, Rachel Stadnitskiy, Sabrina Carmichael, Holger Aulbach, Matthew Hewitt, Xavier De Mollerat Du Jeu, Benedetta di Robilant, Federico Parietti, Jonathan H. Esensten

## Abstract

The production of commercial autologous cell therapies such as chimeric antigen receptor T cells requires complex manual manufacturing processes. Skilled labor costs and challenges in manufacturing scale-out have contributed to high prices for these products. Here, we present a robotic system that uses industry-standard cell therapy manufacturing equipment to automate the steps involved in cell therapy manufacturing. The robotic cluster consists of a robotic arm and customized modules, allowing the robot to manipulate a variety of standard cell therapy instruments and materials such as incubators, bioreactors, and reagent bags. This system enables existing manual manufacturing processes to be rapidly adapted to robotic manufacturing, without having to adopt a completely new technology platform. Proof-of-concept for the robotic cluster’s expansion module was demonstrated by activating and expanding human CD8+ T cells. The robotic cultures showed comparable cell yields, viability, and identity to those manually performed. Such modular robotic solutions may support scale-up and scale-out of cell therapies that are developed using classical manual methods in academic laboratories and biotechnology companies.

## Introduction

Cell and gene therapies are expensive to develop and manufacture, which limits access by patients who may benefit from them [1]. Up to 50% of the manufacturing cost is driven by labor costs and limited availability of skilled labor [2]. Complex manufacturing processes are also difficult to scale up, since manual processes that are cost-effective for small scale production may not be appropriate for large scale commercial manufacturing [3]. The high cost and low throughput of classified cleanroom space is another barrier to development and scale of such therapies [4].

Robotics and automation is a common approach to reducing labor costs in traditional industries such as agriculture [5] and pharmaceutical manufacturing [6]. In autologous cell therapy manufacturing, closed and automated systems such as the Miltenyi Prodigy [7,8] and Lonza Cocoon [9] are widely used as alternatives to traditional manual manufacturing methods. These systems have the advantage of providing an all-in-one solution to cell therapy manufacturing in an instrument that can be operated outside of classified cleanrooms. However, these systems are proprietary and therefore limited in the number and complexity of different manufacturing steps that they can perform. They also require extensive user training and regular user manipulations to add materials and reagents, remove waste, and take samples for quality control analysis. Sterile welding in particular is an error-prone manual process that is used to create sterile connections for closed systems [10].

An alternative approach to such “all in one” manufacturing platforms is a robotic system that can operate industry standard equipment from multiple different vendors already extensively deployed for cell and gene therapy manufacturing. Standard existing equipment is designed for use by human hands; therefore, the robotic system utilizes customized interfaces and equipment workholding to be able to interact with each individual piece of equipment. Examples of such integrated automation approaches include the AUTOSTEM platform for human MSCs [11] and the Stem Cell Factory for induced pluripotent stem cells [12]. Due to their modularity, such systems can rapidly be modified to incorporate new equipment and new techniques as the field evolves. This modularity also allows for processing of multiple different batches in parallel, which is not possible with “all in one” manufacturing platforms.

Here, we describe the performance of a new prototype robotic system designed to operate industry-standard cell therapy manufacturing equipment. We robotically automate the culture of CD8+ T cells using standard G-Rex 10M CS and Xuri Cell Expansion System W25 platforms, in combination with a modified Heracell Vios 250i cell culture incubator. We show that cell yields, phenotype, and viability are comparable between robotic and manual processes. The use of industry-standard equipment in a robotic process can greatly increase the pace of both process development and scale up for novel cell therapy products.

## Materials and Methods

### Robotic Cluster Design

The robotic cluster was designed to enable automated manufacturing of cell therapies. The features of the system are described in Table 1.

**Table 1:**
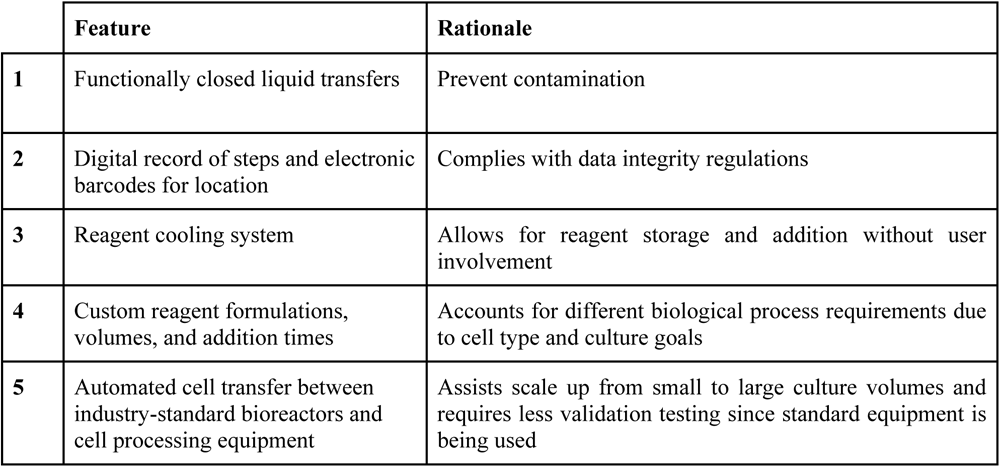
Key features.

|  | Feature | Rationale |
| --- | --- | --- |
| 1 | Functionally closed liquid transfers | Prevent contamination |
| 2 | Digital record of steps and electronic barcodes for location | Complies with data integrity regulations |
| 3 | Reagent cooling system | Allows for reagent storage and addition without user involvement |
| 4 | Custom reagent formulations, volumes, and addition times | Accounts for different biological process requirements due to cell type and culture goals |
| 5 | Automated cell transfer between industry-standard bioreactors and cell processing equipment | Assists scale up from small to large culture volumes and requires less validation testing since standard equipment is being used |

The robotic cluster consists of a central robotic arm surrounded by individual modules that are used to accomplish specific tasks. Each cube-shaped module contains a specific instrument, reagent, or material that, in this proof-of-concept, was used to automate cell expansion (Figure 2a). The robot is controlled by an in-house developed cloud-based software, which records every action and tracks materials through the production process.

#### Interfaces with off-the-shelf equipment

The robotic cluster was designed to use the following off-the-shelf materials and equipment: a G-Rex 10M CS cell culture vessel (Wilson Wolf, US-MN), a Heracell Vios 250i incubator (ThermoFisher Scientific, US-MA), and a Xuri Cell Expansion System W25 (Cytiva, US-MA). The robotic cluster can also handle bags of reagents and syringes. We describe the supporting input and storage sub-modules (Figure 1a and d). Since most off-the-shelf instruments and materials for cell therapy manufacturing are not designed for robotic manipulation, customized cartridges were developed and 3D printed. These cartridges allow the robotic arm to connect tubes, transfer liquids in and out of bags, and move bioreactors that were originally designed for human hands.

**Figure 1:**
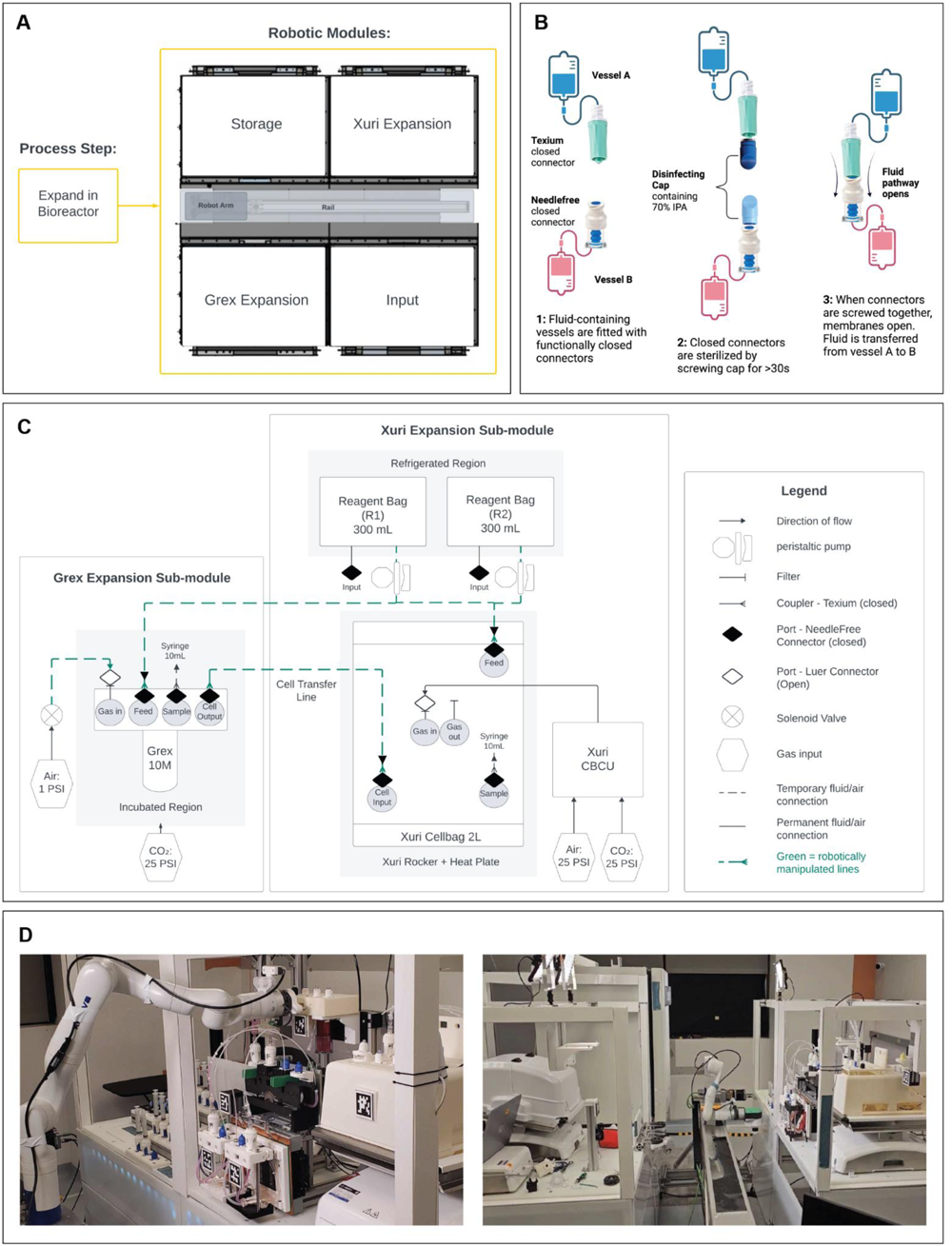
a) outline of the robotic cluster modules used for cell expansion; b) overview of the aseptic disinfection process developed; c) scheme of the G-Rex and Xuri modules indicating robotically manipulated lines; d) photographs of the expansion modules deployed in a CNC lab environment

#### Connections between vessels

Moving of liquids from one vessel to another requires standardized connections between them. Therefore, all consumables used in the process were fitted with tubing extensions containing standard needle free connectors (Texium, BD). These extensions were pre-sterilized using electron beam radiation (Nutek, US-CA) and added to consumables within a BSC. To minimize the risk of microbial contamination during connection of tubes, disinfecting caps containing 70% isopropyl alcohol pads (Merit Medical, US-UT) were used before each connection to robotically sanitize the exposed face of the needle-free connectors (Figure 1b). Disinfecting caps, traditionally used for sanitizing IV lines, may decrease the risk of introducing microorganisms into the sterile fluid path of the tubes [10,13].

#### Cell resuspension and transfer

Resuspending cells and transferring them from one vessel to another is a challenging task for a robotic arm. To transfer cells from the G-Rex culture vessel to the Xuri bag, the robot mixed the G-Rex using an optimized swirling motion to ensure adequate resuspension of cells, and made necessary fluid transfer connections. A compressed air source was then used to transfer the cells to the Xuri culture bag. Compressed air, peristaltic pumps, and syringes operated by the robotic arm are the three methods by which liquids are transferred in the robotic cluster. Wash steps are also included when necessary to ensure adequate cell recovery.

### Reagent storage and addition

Reagents were stored in 300 mL bags with one surface contacting a metal surface that is held at 4°C using the Peltier effect (when electric current flows through the device, heat transfers from one side to the other, leaving one side cooler). The media was separated into two bags labeled R1 (531,000 I.U. of IL-2 diluted in 300 mL cell growth media) and R2 (300 mL cell growth media), dispensed at different ratios to deliver small volumes of IL2 to bioreactors. Reagents R1 and R2 were added (culture day 1, 4, 7, 9, 11) using two concentrically mounted peristaltic pumps (Figure 1c).

### Cells

Human CD8+ T cells isolated from commercially available human leukopheresis collections (StemExpress, US-CA or AllCells, US-CA) using Miltenyi CD8+ microbeads and a Miltenyi Prodigy instrument were used for run 1. Purified human CD8+ T cells were purchased from a commercial source IQ Biosciences and used for runs 2 and 3 (US-CA).

### Reagents

Cell expansion media consisted of CTS optimizer media (Gibco, US-NY) with 5% AB human serum (Access Biologicals, UC-CA) and 1% L-glutamine (Thermo Fisher Scientific, US-CA). Interleukin-2 (IL-2; Clinigen, US-CA) was diluted in phosphate buffered saline (PBS; Thermo Fisher Scientific, US-CA) and added at a final concentration of 300 I.U./mL. Transact (Miltenyi Biotec, US-CA) was used for activation at 10 μL per 1 mL medium.

### *In-vitro* expansion of CD8+ T cells

Stimulation and expansion of human CD8+ T cells was used as a model system for demonstrating the capabilities of the robotic cluster. A robotic culture was performed and compared to a manual control (Figure 2). In brief, cryopreserved CD8+ T cells were thawed, rested overnight in medium, and 15x10^6^ cells were stimulated the next day with TransAct in a G-Rex 10M bioreactor. The G-Rex was kept in a standard cell culture incubator at 37 °C and 5% CO2. The total volume of medium was increased to 90 mL over the first 4 days. On Day 7, the cells were transferred to a Xuri bioreactor (rocking speed 6 RPM, Rocking angle 6°, CO2 5%, air 20%, gas flow rate 0.01 L/min). The total volume was gradually increased to 590 mL. Cultures were terminated on Day 12.

**Figure 2:**
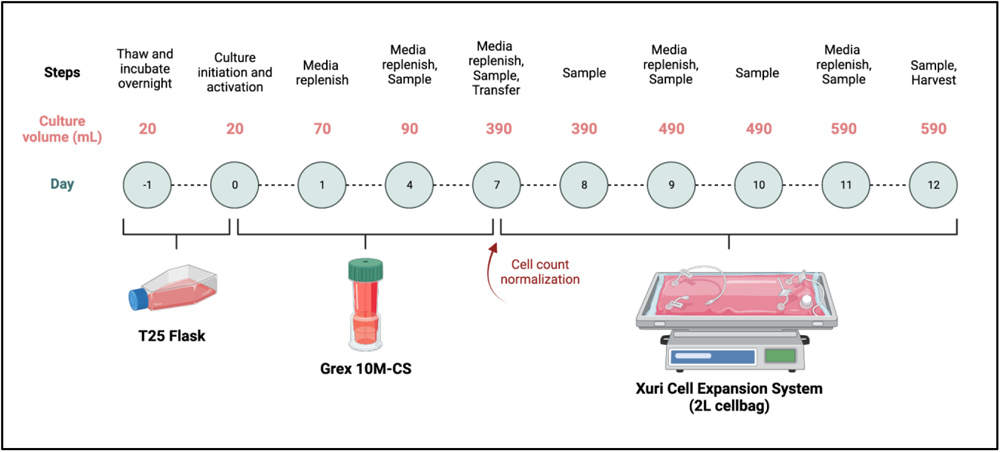
Process steps for manual and automated culturing of CD8+ T cells *versus* process days. Automation started on day 0 post T25 flask thawing/activation.

The same cell expansion was performed by the robotic cell culture, except the G-Rex was placed into the robotic cluster on day 0 after cell seeding and activation. All subsequent manipulations including addition of medium and transfer of the cells from the G-Rex to Xuri were performed by the robotic cluster. Periodic sampling of the culture was performed manually using a serological pipette (G-Rex) or syringe (Xuri) for both robotic and manual conditions. On day 7, the number of cells transferred from the G-Rex to the Xuri was equalized between the robotic and manual conditions. An additional control culture was performed in traditional plastic flasks with periodic media additions (Supplementary figure S1).

### Cell characterization

#### Cell Count and Viability

Cell count and viability analysis was performed on 200 uL of sample using an NC200 Nucleocounter (Chemometec, US-CA) and Via2 cassettes. Calculations for cell yield and doubling time are referenced in the supplementary section.

#### Flow Cytometry

Cryopreserved samples were thawed, washed with PBS, and resuspended in BD Stain Buffer (Catalog Number 554656). Fluorochrome-conjugated antibodies used for staining are listed in Table S1. Fluorescent minus one (FMO) controls, and single-color controls using UltraComp Beads were prepared. Flow cytometry was performed by the UCSF Flow core on a Sony ID7000 Spectral Cell Analyzer. Data was analyzed in FlowJo (BD).

#### Bulk RNA Sequencing

Bulk RNA sequencing was performed on T cells cryopreserved on day 11 or 12 by Novogene (US-CA), as previously described [14]. After thawing, RNA was extracted and RNA degradation and contamination was monitored on 1% agarose gels. RNA purity was checked using the NanoPhotometer^®^ spectrophotometer (IMPLEN, US-CA). RNA integrity and quantitation were assessed using the RNA Nano 6000 Assay Kit of the Bioanalyzer 2100 system (Agilent Technologies, CA, USA). A total amount of 1 μg RNA per sample was used as input material for the RNA sample preparations. Sequencing libraries were generated using NEBNext® UltraTM RNA Library Prep Kit for Illumina® (NEB, US-CA) following manufacturer’s recommendations and index codes were added to attribute sequences to each sample. At last, PCR products were purified (AMPure XP system) and library quality was assessed on the Agilent Bioanalyzer 2100 system. The clustering of the index-coded samples was performed on an Illumina Novaseq sequencer according to the manufacturer’s instructions. After cluster generation, the libraries were sequenced on the same machine and paired-end reads were generated. Mapping was performed by histat2 (2.0.5), quantification by featureCounts and differential analyses using DESeq2 (1.20.0) and edgR (3.22.5) |log2(FoldChange)| >= 1 & p <= 0.05. The NovoMagic software was used for visualization and other analyses. Gene enrichment analyses were performed using the GSEA software (broad Institute). The Broad Institute gene set database was used as a reference (h.all.v2023.2.Hs.symbols.gmt and c7.immunesigdb.v2023.2.Hs.symbols.gmt were used). The annotation platform was set to Ensembl (Human_ENSEMBL_Gene_ID_MSigDB.v7.2.chip).

#### Aseptic Process Simulation

A subset of robotic operations, including reagent addition, sampling, and cell transfer from G-Rex to Xuri, were simulated over the course of 12 days using Tryptic Soy Broth (TSB; Sigma Aldric, US-CA). The final Xuri bag was tested for turbidity after a 14 day incubation period, with a positive control experiment performed in parallel (*i.e.* inoculation of the bag with <100 CFU of USP indicator organism).

#### BacT sterility test

At the end of the process, samples from the manual and robotic conditions were collected for sterility analysis in accordance with USP <71> in Bactec anaerobic and aerobic bottles or in 15 mL conicals for fungal/gram stain (Becton Dickinson, US-NJ). Bactec bottles were incubated and checked for bacterial growth for 14 days.

### Statistical Analysis

The statistical significance between manual versus robotic conditions for yield, viability, day 7 fold expansion, day 12 fold expansion, and surface marker expression (PD1+, LAG3+, TIM3+) was assessed using Prism (Graphpad Prism, US-CA) paired T tests with significant P <0.01. RNA sequencing results were analyzed for differential expression with the DESeq2R package (1.20.0); resulting p-values were controlled for false discovery rate using Benjamini and Hochberg’s approach. Differentially expressed genes had an adjusted p-value of <=0.05 found by DESeq2.

## Results

To assess the robotic system’s ability to culture T cells with similar performance to a manual system, manual and robotic conditions were tested in parallel. A culture of 15x10^6^ CD8+ primary human T cells was stimulated in a G-Rex 10M-CS on day 0 and placed into a cell culture incubator. The cells were transferred on day 7 to a Xuri Cell Expansion System W25. The cultures were maintained in the Xuri through day 12 and were sampled at regular intervals over the culture period. Overall results from parallel robotically and manually performed cell cultures are summarized in Figure 1. Total cell yield and cell viability on days 7 and 12 were not significantly different between robotic and manual processes (Figure 3A). We confirmed that the temperature inside the Xuri culture bag was similar between the two conditions, since the robotic condition used a customized bag scaffold that could disrupt normal heat flux (Figure S1). The efficiency of the transfer process of the culture from the G-Rex to the Xuri bioreactor on day 7 was evaluated by measured total cell number before and after the transfer (Figure 3B). There was no significant difference in percentage of cells transferred between the robotic (102%) and manual (106%) conditions (Figure 3B, right panel). When cell viability was measured both before and immediately after transfer on day 7 from G-Rex to Xuri, there was a trend toward lower viability in the robotic condition (Figure 3C). However, the difference was not statistically significant and the cultures recovered high viability by the end of the culture period. Additional comparison of the manual and robotic conditions is presented in Table S3. Bacterial culture was performed on samples from the cell culture collected on day 12 of each run to confirm sterility. In one of the three manual T cell expansions, the cultures turned positive at 21 hours after the beginning of culture. *S. epidermidis* was identified in two separate culture bottles. None of the samples from the robotic cell expansions grew bacteria.

**Figure 3.**
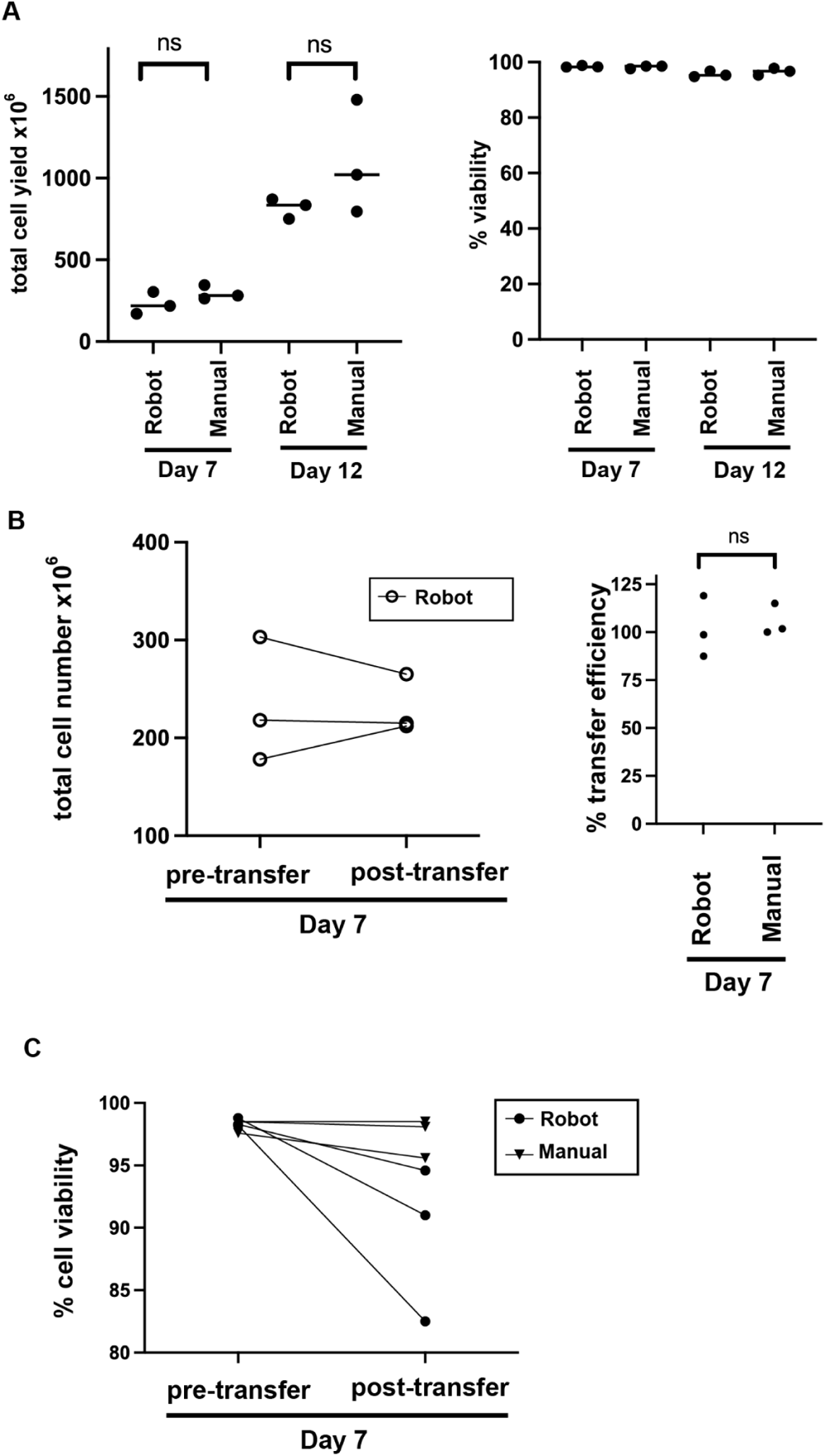
Comparison of expansion of stimulated primary human CD8+ T cells using the robotic system and an analogous manual process. A) Total cell yield and viability at day 7 and day 12 of the culture. B) Cell numbers immediately before and after transfer of cells from the G-Rex to the Xuri on Day 7, with the transfer efficiency. Post-transfer cell number for manual process was matched to the number of cells transferred by the concurrent robotic process, so post-transfer cell number is not informative for the manual process. C) Cell viability before and after cell transfer at Day 7.

### Optimization of cell resuspension and transfer via robot

The transfer of cells from the G-Rex 10M CS to the Xuri Cell Expansion System W25 requires resuspension of cells which have settled to the bottom of the G-Rex culture vessel, followed by transfer of the culture volume to the Xuri. Although such resuspension and transfer is simple using a manual pipette, resuspension and transfer of cells via the G-Rex tubing is more challenging for a robot, because cells adhere to the G-Rex mesh. Therefore, we explored two different approaches to resuspend cells in the G-Rex vessel prior to transfer by either tilting or swirling motions, which are compatible with single-joint robotic manipulation. A total of 3.0 x 10^6^ primary human T cells in 90 mL of culture medium were allowed to settle for >2 hours and then the cells were either tilted manually 10 times at a low angle or swirled with a top pivot 10 times. Following this step, the cells were transferred out of the G-Rex via a syringe connected to the G-Rex harvest port to a second vessel. The number of cells transferred was counted, and then 2 washes were performed with 20 mL each of media. Resuspension methods (tilt or swirl) were used again after each wash step, before transferring. Additional cell counts were taken after each of the washes. These resuspension procedures were repeated 3 times with intervening addition of wash media, and both methods demonstrated >95% efficiency of cell transfer after 3 rounds of resuspension and transfer (Figure 4, left and middle panels). The “tilt” method was then tested using the robotic cluster instead of human hands. There was >98% transfer of cells out of the G-Rex using this method on the robotic cluster (Figure 4, right panel).

**Figure 4:**
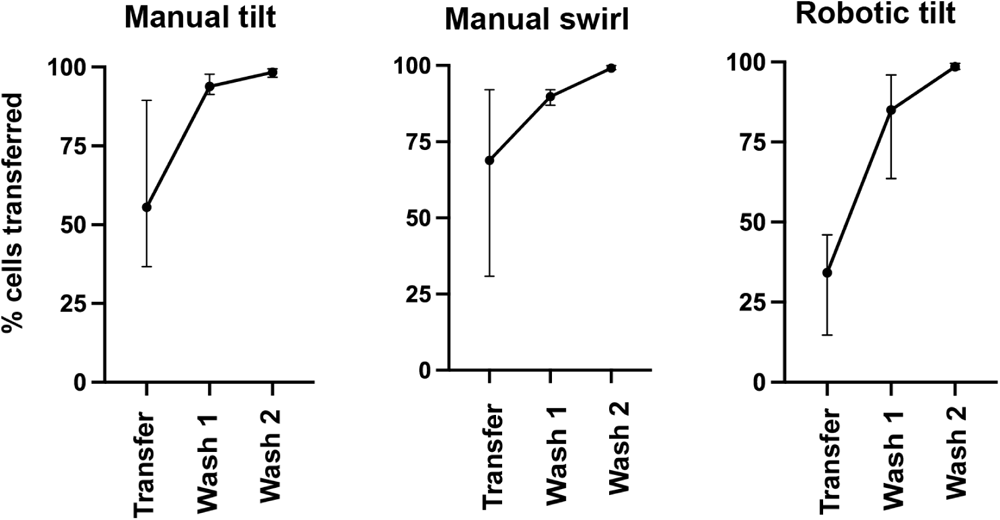
Optimization of cell transfer from G-Rex using different methods for resuspension of cells. 3.0 x 10^6^ cells in 90 mL were transferred from a G-Rex after manual resuspension via tilting (left) or swirling (center). The tilting or swirling was repeated before the initial transfer and with each wash step (20 mL used per wash). The tilting method was adapted to a robotic process and the efficiency of cell transfer was confirmed (right). Error bars indicate range. Data are from 3 independent experiments.

### Sterility of Robotic Processing

To confirm the ability of the robotic system to maintain sterility during cell manipulations and culture, we performed aseptic process simulations with TSB medium (n=1) as described above. No bacterial growth was observed in the media at the end of the process in which the whole volume of culture medium was incubated for 14 days at 37 °C. This process was carried out in standard (non-cleanroom) laboratory space. Analysis of the levels of viable and non-viable particles in the environment of the robotic cluster is summarized in Table S2, which demonstrates that the robotic cluster maintains sterility of the culture even in a non-classified environment.

### Comparison of cell phenotype

To explore potential differences in cell phenotype between cultures handled manually versus those handled by the robotic cluster, we performed bulk RNA sequencing on specimens from day 11 of the cultures. A hierarchical cluster analysis of gene expression was performed for the three parallel runs (Figure 5A) which showed clustering by run and not by robotic vs manual handling. Analysis of differentially regulated genes by volcano plot of manual vs robotic condition (Figure 5B) showed a small number of differentially expressed genes: 128 genes, 0.45% of the total analyzed. Comparison of differentially regulated genes between paired manual and robotic runs (Figure 5C) showed limited overlap in genes that were differentially regulated between conditions. None of the 9 genes found to be differentially regulated in all 3 replicates (Figure 5D) related to T cell expansion, exhaustion or senescence. Expression of a specific subset of genes involved with differentiation, exhaustion, and proliferation was compared and found to be statistically indistinguishable between the manual and robotic conditions (Figure 5E).

**Figure 5:**
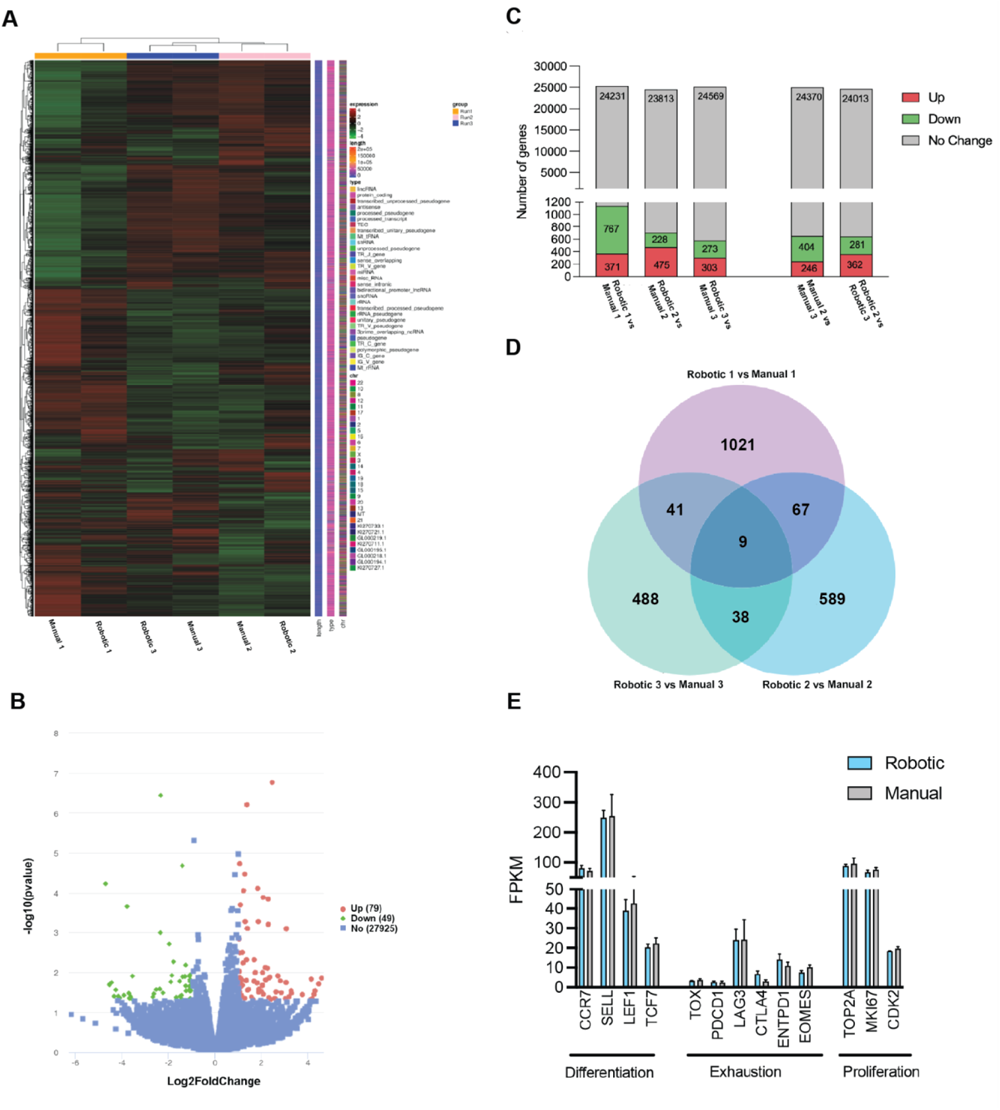
RNA sequencing results. Differentially expressed genes calculated using P value <0.05 and |log2FoldChange|>1. a) hierarchical clustering for robotic *versus* manual conditions (n=3); b) volcano deg plot for all manual *versus* robotic conditions (n=3); c) up and down regulated genes for each manual *versus* robotic run, manual run 2 *versus* manual run 3, and robotic run 2 *versus* robotic run 3; d) venn diagram of differentially expressed genes for all manual *versus* robotic conditions (n=3); e) Differentiation, exhaustion, and proliferation genes for all robotic *versus* manual conditions (n=3), bars indicate standard deviations.

Flow cytometry analysis performed on samples from days 7 and 11 of the cultures showed >95% expression of CD3+ and CD8+ markers (Figure 6A). Expression of exhaustion markers PD1, LAG3, and TIM3 was similar between manual and robotic conditions (Figure 6B). Subset analysis using CD45RO and CCR7 showed a preponderance of CD45RO+CCR7+ Tcm cells in both manual and robotic cultures.

**Figure 6:**
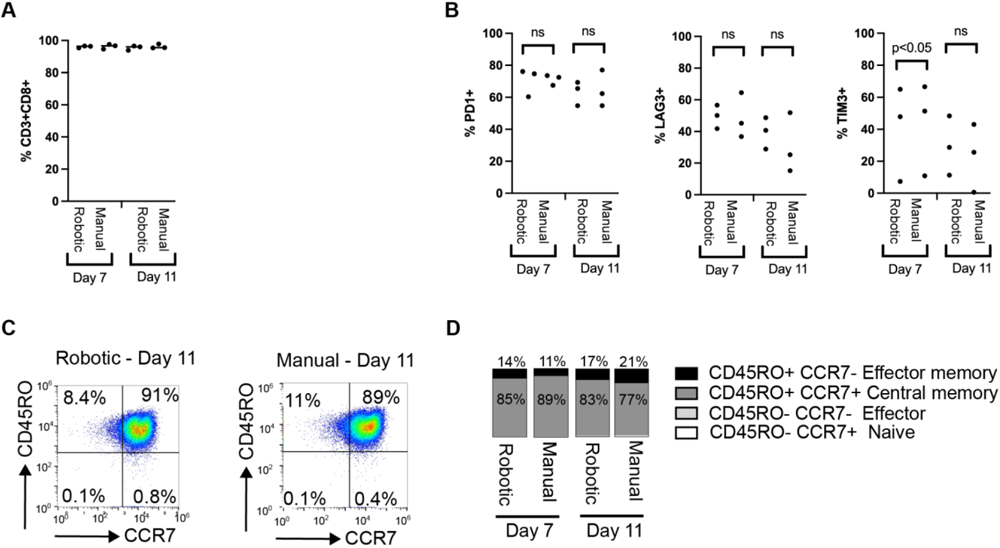
Flow cytometry analysis of robotic *versus* manual cultures at day 7 and 11. A) Expression of CD3+CD8+ in cultures. B) Expression of exhaustion markers PD1, LAG3, and TIM3. C) Gating strategy to identify T cell subsets. D) Average of T cell subsets in each condition (n=3).

## Discussion

We have presented the design and performance data for a functionally closed robotic cluster with core functionalities for culturing human T cells in industry-standard bioreactors. The robotic cluster stores and adds medium, transfers cells, and maintains bioreactor cultures for 12 days without direct human intervention (Figure 2). The robotic cluster is designed to be modular so that it can incorporate any industry-standard equipment into its workflow. Each piece of equipment integrated into the robotic cluster is controlled by a cloud-based control system that interfaces directly with the equipment’s application programming interface (API). We presented data from expansion of primary human CD8+ T cells (n=3) and demonstrated comparable results between the process as performed by human hands and by the robotic system (Figure 3). Future work will focus on expanding the number of manipulations that the robotic cluster can perform, such as automated density gradient centrifugation and T cell subset isolation at the beginning of a process, electroporation for genetic modification in the middle of the process, and formulation, fill, and finish at the end. Since there are already industry-standard instruments for each of these steps, it will be possible to rapidly integrate these and other processes into the robotic cluster.

Importantly, the robotic cluster requires customized cartridges to be able to manipulate a wide variety of standard consumables such as syringes, plastic culture vessels, and culture bags. Each cartridge is designed to maintain the as-designed performance of the consumables while making the consumables amenable to robotic manipulation. In our development, we encountered a problem in which a customized cartridge interfered with temperature regulation in the Xuri bioreactor. However this problem was overcome through iterative improvements to the design of the cartridge. In addition, the manufacture of pre-filled bags, vials, or syringes with commonly used consumables will be important for future development, since the robotic cluster has limited ability to handle small volumes required for some materials (such as cytokines or activation reagents).

There are some important differences between manual and robotic manipulations in the presented data: the robotic arm moves more slowly than human hands, so the cells in the robotic condition spend more time outside the controlled temperature of the incubator during transfer steps (0.5-1.5 hours) compared to less than 15 minutes for manual transfers. The resulting variability during these transfer steps could have measurable biological impacts on the cells, although no such differences were seen in our phenotyping data by bulk RNA sequencing (Figure 5) or flow cytometry (Figure 6). Similarly, we show that a robotic approach to resuspending cells in a Grex bioreactor using a tilting motion has comparable efficiency to manual tilting (Figure 4).

Sterility of the robotic process is maintained through the use of multiple-use needleless connectors, which are sanitized with alcohol-containing disinfecting caps before each use. The development of on demand aseptic connections provides advantages compared to sterile welding since it is easier for a robot to perform and allows for multiple connections to be made and removed from the same vessel. Future development of this system will include support for different equipment for additional process steps (*e.g.* electroporation, transduction, separation, freeze, thaw) and the ability to work with more bioreactors in parallel, all unified by overarching hardware/software. In the data presented above, we show that the sterility of the culture is maintained by the robotic process, while in one case the control manual process resulted in a positive bacterial culture from an end-of-process cell sample. The recovered organism (*S. epidermidis*) is found on human skin and supports the conclusion that removing human hands from manufacturing processes can help to prevent microbial contamination.

In summary, a robotic system with high modularity and flexibility can be leveraged to operate a range of standard equipment, automate a variety of process steps, and allow for multiple products cultured in parallel. This technology can be used for parameter exploration during process development since it can easily run multiple processes in parallel. It can also be used to scale up clinical manufacturing of autologous cell therapies in a small footprint. Therefore, future development of robotic systems for cell therapy manufacturing has the potential to increase quality while decreasing labor costs and speeding up scale-up.

## Conclusions

A prototype robotic cluster for manufacturing cell therapy products demonstrated comparable cell yields, viability, and identity when compared to manually cultured CD8+ cells. The use of industry-standard equipment in the robotic cluster may support accelerated manufacturing scale-up. Further development of robotic approaches to cell therapy manufacturing has the potential to increase quality, decrease labor costs, and increase manufacturing throughput.

## Supplementary Material

Figure S1. Temperature study with customized Xuri bag cover

**Figure S1:**
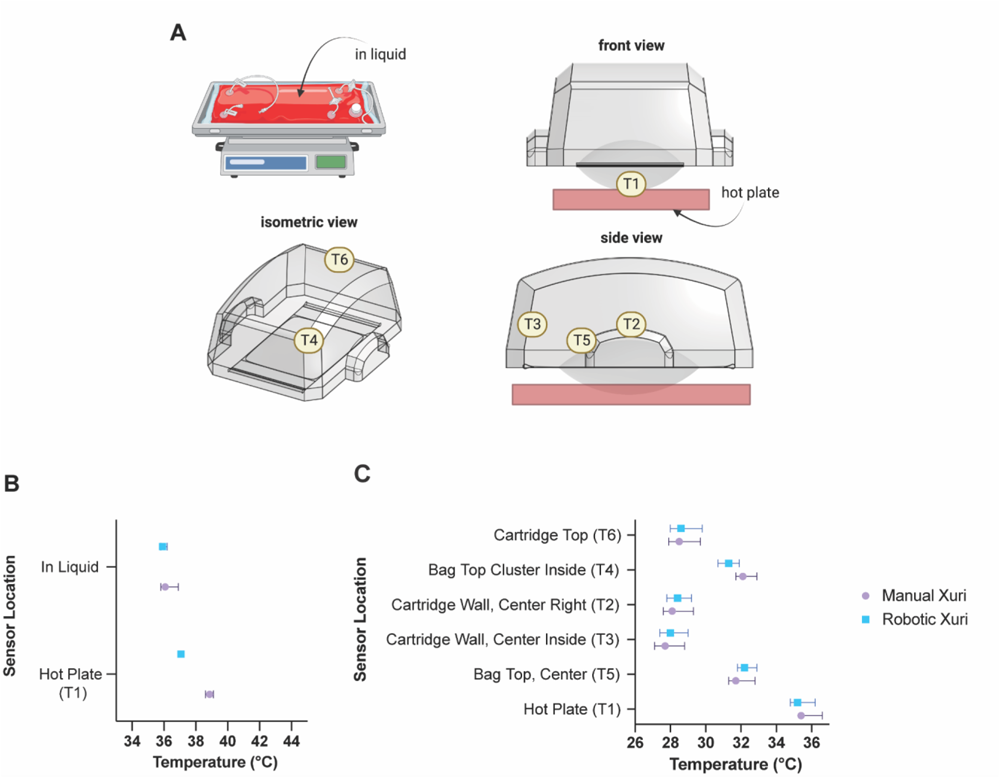
Xuri temperature study results. a) outline of the temperature sensor locations, b) comparison of in-liquid *versus* hotplate temperatures for manual *versus* robotic Xuris and c) air temperature at different locations within the manual versus robotic Xuri bag covers. Bars indicate standard deviations.

Since a new 3D printed cover had to be created for the robotic arm to manipulate the Xuri culture bag, thermal equivalence studies were conducted to confirm that Xuri temperature within the robotic system exactly matched that of the manual Xuri. Two Xuri bioreactors were compared: one with the standard bag and plastic bag cover, and one with the customized robotic cover. For both setups, 350 mL of distilled water was added to 2 separate Xuri bags and allowed to equilibrate. Temperature measurements were taken over the course of 24 hours (probes with +/-1 °C accuracy, EXTECH TM500, US-MA) and sampled every minute. Temperature evaluation was assessed in different locations, even including air, in liquid and hot plate measurements (S1).

Figure S2. Supplementary RNA Sequencing

**Figure S2:**
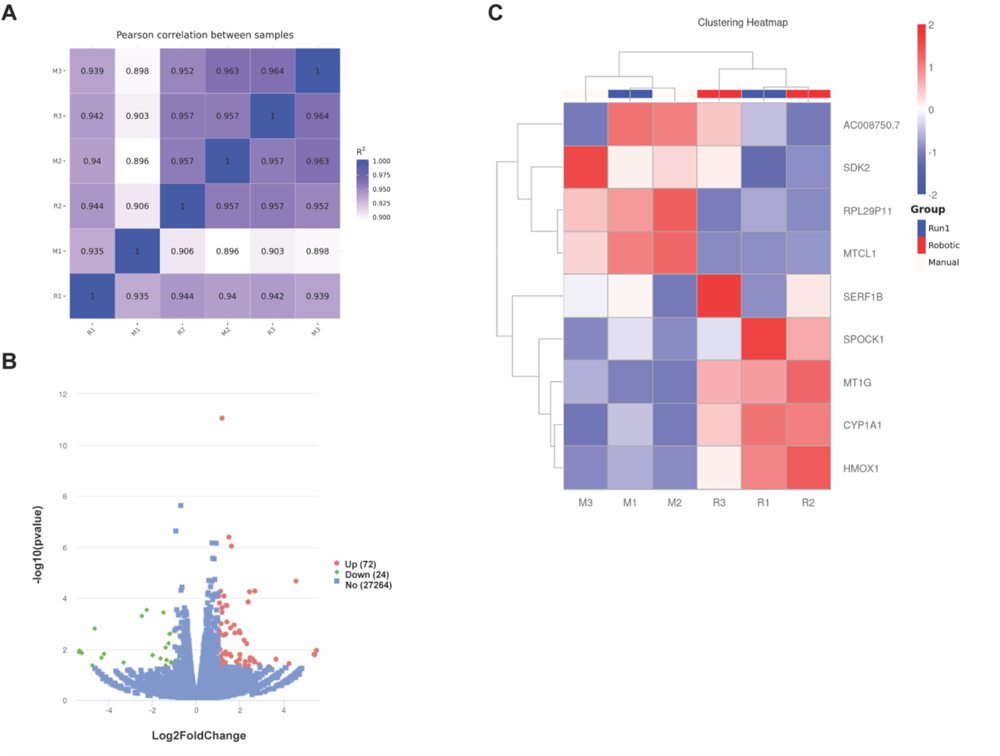
RNA sequencing supplemental results. Differentially expressed genes calculated using P value <0.05 and |log2FoldChange|>1. a) Pearson correlation between robotic *versus* manual conditions (n=3); b) volcano deg plot excluding run 1 due to contamination (n=2); c) clustering heat map (n=3).

**Table S1:** Flow panel antibodies.

|  | Antibody | Vendor | Catalog number |
| --- | --- | --- | --- |
| Panel 1 | CD3 FITC | BD | 561802 |
|  | TIM3 BB700 | BD | 746178 |
|  | CD8 APC H7 | BD | 560273 |
|  | LAG3 BV421 | BD | 565721 |
|  | Live Dead Aqua Blue | Thermofisher | L34965 |
|  | PD1 BV786 | BD | 563789 |
|  | CD4 BUV395 | BD | 563550 |
| Panel 2 | CD45RA FITC | BD | 561882 |
|  | CXCR3 BB700 | BD | 566533 |
|  | CCR7 APC | BD | 566762 |
|  | CD8 APC H7 | BD | 560273 |
|  | CD49D BV421 | BD | 565277 |
|  | Live Dead Aqua Blue | Thermofisher | L34965 |
|  | CD45RO BV786 | BD | 564290 |
|  | CD27 BUV395 | BD | 563816 |
|  | CD4 BUV805 | BD | 612888 |

**Table S2:** results of aseptic simulation tests.

| **Environmental monitoring: active particle counts** |  |  |  |
| --- | --- | --- | --- |
| **Sample Location** | **Avg PPCM count (um) >0.5** | **Avg PPCM count (um) >5.0** | **Location Type** |
| 1 - robotic rail bottom | 202380 | 3120 | standard research lab |
| 2 - input module | 203260 | 2460 | standard research lab |
| 3 - robotic rail top | 389500 | 1940 | standard research lab |
| 4 - storage module | 200540 | 1540 | standard research lab |

| **Environmental monitoring: viable colony counts** |  |  |  |
| --- | --- | --- | --- |
| **Sample Location** | **Settling plate CFU count** | **Contact plate CFU count** | **Location Type** |
| 1 - Xuri sub-module | 2 | 19 | standard research lab |
| 2 - Storage sub-module | 6 | 6 | standard research lab |
| 3 - G-Rex sub-module | 2 | 0 | standard research lab |
| 4 - Input module | 12 | 7 | standard research lab |

| **Tryptic Soy Broth Incubation of APS** |  |
| --- | --- |
| **Condition** | **Final Result** |
| Pre-wash bag (aseptic assembly control) | No growth to 14 days |
| Xuri product bag | No growth to 14 days |

**Table S3:** cell expansion results.

|  | **Parameter** | **Manual Condition**<br>**(n=3)** | **Robotic Condition**<br>**(n=3)** | **Statistics**<br>**(paired T test)** |
| --- | --- | --- | --- | --- |
| **G-Rex** | Seeding number ($\times 10^7$ cells) | 1.5 | 1.5 | n/a |
| | Seeding density ($\times 10^6$ cells/mL) | 0.75 | 0.75 | n/a |
|  | T cell doubling level | 4.31 | 3.92 | 0.13 (ns) |
|  | T cell doubling time (days) | 3.02 | 3.34 | 0.16 (ns) |
| | Pre-transfer viability (%) | $98.2 \pm 0.49$ | $98.4 \pm 0.71$ | 0.68 (ns) |
| | Pre-transfer live cells ($\times 10^8$ cells/mL) | $2.99 \pm 0.38$ | $2.33 \pm 0.58$ | 0.096 (ns) |
|  | G-Rex-Xuri cells transferred (%) | 101.86% | 105.73% | n/a |
|  | Total cumulative medium (mL) | 90 | 90 | n/a |
| **Xuri** | Seeding number ($\times 10^8$ cells) | $2.33 \pm 0.45$ | $2.31 \pm 0.30$ | n/a |
| | Seeding density ($\times 10^6$ cells/mL) | 0.8 | 0.8 | n/a |
|  | T cell doubling level | 2.36 | 1.99 | 0.10 (ns) |
|  | T cell doubling time (days) | 5.53 | 6.60 | 0.11 (ns) |
| | Harvest viability (%) | $96.6 \pm 1.06$ | $95.6 \pm 1.77$ | 0.21 (ns) |
| | Total live cells ($\times 10^9$ cells) | $1.21 \pm 0.25$ | $0.91 \pm 0.051$ | 0.28 (ns) |
|  | Total cumulative medium (mL) | 590 | 590 | n/a |

### Yield and doubling calculations

Cell yields for robotic *versus* manual experiments were determined by multiplying cell concentration by total media volume, without subtracting volumes used for sampling and cryopreservation, which were consistent between manual and robotic conditions (figure 3, table S3). Cell doubling level (Eq. 1) and doubling time (Eq 2.) were calculated:

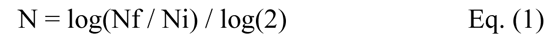

N: doubling level

Nf: final number of cells in the population

Ni: initial number of cells in the population

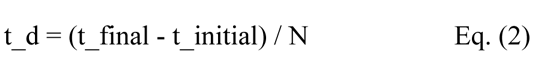

t_d: doubling time

t_final: final time

t_initial: initial time

N: doubling level

